# Biophysical Characterization and Interaction study of WhiB6 Protein of *Mycobacterium tuberculosis* with Nucleic Acid

**DOI:** 10.1101/2023.03.01.530725

**Authors:** Sonam Kumari, Ruchi Singh, Teena, Soumik Siddhanta, Shashank Deep

**Author notes:** **To whom correspondence should be addressed.** Shashank Deep, Professor, Department of Chemistry, Indian Institute of Technology, Delhi (IIT Delhi), Hauz Khas, New Delhi – 110016, INDIA, (O).

## Abstract

Tuberculosis is an intractable disease because of the peculiar nature of the virulent properties of *Mycobacterium tuberculosis*. The WhiB6 protein, a transcriptional regulator, plays a crucial role in the virulence systems of *Mtb*. It regulates the expression of genes essential for the virulence pathways by binding to their promoter region; *espA* is one such gene. Herein, we have used biophysical methods, including steady-state intrinsic fluorescence spectroscopy, circular dichroism spectroscopy, Isothermal titration calorimetry (ITC), and Surface-Enhanced Raman Spectroscopy (SERS) to understand the interaction of WhiB6 protein with *espA* promoter DNA. For the first time, we report the conformational details and biophysical parameters related to the WhiB6-*espA* promoter DNA interaction. WhiB6 binds the DNA with moderate affinity, as revealed by ITC. CD and SERS studies suggest subtle perturbation in the secondary conformation of the protein on binding to the DNA. SERS provided detailed structural insights into the WhiB6 protein and the amino acids involved in the interaction, which could be harnessed to find suitable inhibitors of the protein-DNA interaction. Preventing the binding of WhiB6 with promoter DNA of the virulence genes can hinder the functioning of *Mtb* and hence can act as an effective therapeutic intervention for tuberculosis.

## INTRODUCTION

Tuberculosis (TB) is one of the deadliest infectious diseases worldwide. In 2020, World Health Organization (WHO) reported approximately 1.5 million mortalities due to TB. According to the WHO Global TB report 2021, TB deaths rose for the first time in more than a decade due to the COVID-19 pandemic. A plethora of research is going on to eradicate this disease. The major challenge faced in current TB research is understanding the erratic nature of *Mycobacterium tuberculosis* (*Mtb*), the causative agent of TB. *Mtb* is an obligate aerobe, a type of Actinomycetales bacteria of the Mycobacterium family and Mycobacterium genus. Many fundamental questions about the physiology, molecular pathogenesis, and mechanism used by *Mtb* for its survival during early infection and persistence in the host cell are still unexplored. *Mtb* possesses a distinctive ability to survive under hostile conditions it encounters during infection (1). It maintains the tendency to adapt to different environmental conditions within the host organism. The mechanism used by the bacterium to evade its host’s defenses to survive the vigorous host immune response remains poorly understood. It also possesses a remarkable feature of remaining in a non-replicating persistent state inside the host, emerging from the persistent state, and causing active disease. The modulation of its environment and the ability to enter a dormant phase are the hallmarks of *Mtb*.

*Mtb* utilizes extensive gene regulations to enter into and emerge from the persistent state. Understanding these gene programming and detailed examination of the genes responsible for responding under stress conditions can lay the foundation for discovering promising drugs for TB. In addition to genes, proteins of any organism play a vital role in most biological processes; they can be an attractive target for diagnostics and drug delivery. Proteome studies of *Mtb* have given insights into the role of some proteins, but numerous proteome aspects remain unexplored (1). Genes responsible for responding under stress conditions require detailed examination as well.

An increase in the cases of multidrug resistance (MDR) and extensive drug resistance (XDR) has made it a dire need to look for new TB drugs. WhiB proteins are essential transcription regulators of *Mtb*; they regulate the expression of several crucial genes of *Mtb* by binding to their promoter DNA. *whiB* belongs to the family of *whi* genes; they are the founding member of the WbI family of proteins that carry an iron-sulfur cluster [4Fe-4S] (2). *whi* genes are present exclusively in *Actinomycetes*, such as *Mycobacterium* and *Streptomyces spp*., and are absent from all other organisms studied so far (3). WhiB proteins do not show homology with human proteins. Seven *whiB* genes are known to be present in *Mycobacterium*. They all possess three sequence motifs: four conserved cysteines (C-X19-36-C-X-X-C-X5-7-C) at an N-terminal domain that coordinate an iron-sulfur (Fe-S) cluster (3), conserved G(V/I)WGG turn and basic amino acids at C-terminal. The C-terminal basic amino acids and the tryptophan region form a helix-turn-helix-like structure that is predicted to be involved in DNA binding (4, 5). Their involvement in the crucial functions of *Mtb* has made them an important drug target. Before choosing it as a drug target, there is a sore need to understand these proteins’ precise functions thoroughly. They are involved in cell division (WhiB2), maintenance of redox homeostasis (WhiB3, WhiB4, and WhiB7), virulence (WhiB3, WhiB4, WhiB5, and WhiB6), regulation of protein secretion (WhiB1, WhiB5, and WhiB6), antibiotic resistance (WhiB7) and reactivation from the dormant state (WhiB1 and WhiB5) (6–12). Very few biochemical properties of the WhiB proteins are known; their function and numerous other aspects are still in their infancy. Therefore, there arises a need to investigate the structure and function of WhiB proteins.

The whiB6 protein of the *whi* gene family regulates *Mtb* virulence. It is located adjacent to the ESX-1 gene locus; regulates the expression of ESAT-6 (early secretory antigenic target) protein family secretion (ESX) systems; it is also known as type VII secretion systems (13). ESAT-6 (secreted by ESX-1) is a vital virulence factor of *Mtb*; its inactivation severely affects the virulence tendency of *Mycobacteria* (12). Since WhiB6 regulates the expression of the ESX system’s genes, thus it is pivotal for *Mtb* virulence. The expression of *whiB6* is correlated with the *esxA* (*rv3875*) (codes for ESAT-6). It has been reported that downregulation of *whiB6* by a single nucleotide insertion in its promoter region leads to a decrease in secretion and production of ESAT-6; also, expression of ESX-1 associated genes (*pe35, espE, esxA*, and *esxB*) is reduced in *whiB6*/Rv3862c mutated strains (14). WhiB6 regulates the expression of genes by binding to their promoter region. It interacts with the promoter region of *espA* and ESX-1 genes *pe35* and *espB* (15, 16). These genes play a crucial role in the virulence of *Mtb*; they are required for ESAT-6 secretion (17, 18). After secretion EspA forms dimers by disulfide bond formation; this bond is essential for maintaining cell wall stability and functioning of the complete ESX-1 system (19); *espA* is induced under osmotic stress, so it is presumably crucial for *Mtb* to survive under osmotic stress. Furthermore, WhiB6 plays a pivotal role in aerobic and anaerobic metabolism and cell division (14). Although it is known that WhiB6 binds to DNA, any further information regarding the binding remains obscure.

As mentioned above, WhiB6 regulates crucial genes of *Mtb* by binding to their promoter DNA; transcription of the *whiB6* gene itself is also regulated by other regulators. Proteins belonging to the OmpR/PhoB subfamily, which is the largest of the response regulators (20), regulate whiB genes: PhoP proteins that play a pivotal role in the virulence of *Mtb* positively regulate transcription of *whiB6* by binding to the direct-repeat motifs of its promoter gene (in clinical *Mtb* strains but not in the standard laboratory strain H37Rv) (21). Besides involvement in virulence, WhiB6 exhibits a significant degree of stress modulation in *Mtb*. It shows strong upregulation in response to a wide variety of stress conditions. Exposure to acidic conditions (pH 4.5) upregulates *Mtb whiB6*. Along with biochemical analyses of the WhiB6 gene products, this information will be necessary for understanding the biology of this novel protein of *Mycobacteria* (22).

The cysteine residues of WhiB6 coordinate the iron-sulfur [4Fe-4S] cluster (23). It exists in the holo-WhiB6 form when the (FeS) cluster is coordinated by the cysteine residues, and once the cluster is lost, it transforms into the active apo-WhiB6 form. Moreover, it has been reported that on the removal of the iron-sulfur cluster, the cysteine-thiols of all the WhiB proteins undergo oxidation and form two intramolecular disulfide bonds (23). WhiB proteins exhibit DNA binding properties and show protein disulfide reductase activity (except WhiB2) in the Apo-from. The reason for the gain of disulfide reductase activity in the apo form has been explained in detail in a previous study (Alam et al. (2008)). The cysteine-thiols in the holo-form are not accessible for electron flow and disulfide exchange when they remain ligated to the cluster. Upon exposure to the oxidizing conditions, the iron-sulfur cluster disintegrates, leading to conformational change accompanied by the gain of disulfide reductase activity (23). It can be inferred that on experiencing redox stress, loss of the cluster of the WhiB proteins is accompanied by a gain in DNA-binding and protein disulfide reductase activities.

WhiB6 is a redox-sensitive protein. Its conformation and function alter under oxidizing and reducing conditions (23). Oxidation and reduction of the cysteine residues are enough to change the protein conformation, abolish or promote interaction with DNA, and activate or repress transcription. Chawla et al. (2012) reported that oxidation of cysteine thiols of apo-WhiB4 induces DNA binding and represses transcription of WhiB4, whereas reduction reverses the effects (24). The Redox state of apo-WhiB3 significantly affects its DNA binding efficiency. Reduction of the apo-WhiB3 cysteine thiols abolishes its DNA binding capabilities, whereas oxidation restores it (25). According to the previously reported studies, the apo form of WhiB proteins (WhiB1, WhiB2, WhiB3, and WhiB4) bind to DNA more strongly than the holo form, and in the apo-oxidized state, they bind DNA with higher affinity than the reduced state (8, 11, 26, 27). Hence, the WhiB6-DNA interaction studies has been investigated under both oxidized (Apo WhiB6-(S)_2_) and reduced (Apo WhiB6-(SH)_2_) conditions.

In this study, for the first time, we report the biophysical characterization and reductase activity of the recombinant WhiB6/Rv3862c protein of *Mtb* and utilized ITC, circular dichroism spectroscopy, steady-state fluorescence spectroscopy, and surface-enhanced Raman spectroscopy (SERS) to corroborate the interaction between WhiB6 and the promoter DNA of a virulence gene (*espA*) experimentally. As above-mentioned, *espA* is regulated by WhiB6; hence, the interaction between WhiB6 and the promoter DNA of the *espA* gene has been investigated using the aforementioned biophysical techniques. Since no anti-virulence drugs have entered clinical development yet, this information related to interaction will be advantageous in designing a drug that could obstruct the expression of virulence genes regulated by the WhiB6 protein.

## MATERIALS AND METHODS

### Materials

Luria agar and Luria Bertani broth for bacterial culture were procured from Himedia. 8-anilino-1-naphthalene sulfonate (ANS) was purchased from Sigma-Aldrich. Ni-NTA resin was obtained from Qiagen. Tris buffer and sodium chloride (NaCl) were obtained from Merck. Tris(2-carboxyethyl)phosphine hydrochloride (TCEP.HCl), Dithiothreitol (DTT), and imidazole were obtained from SRL, India. All the reagents used were of analytical grade.

### Expression and Purification of WhiB6 protein

pET21a Plasmid encoding *Mtb* whiB6/Rv3862c gene fused with His_6_ tag was transformed in BL21(DE3), carrying a copy of the T7 RNA polymerase under the control of the IPTG-inducible lacUV5 promoter competent cells. After 4 hours of induction with IPTG at 37 °C, the bacterial culture was harvested by centrifugation at 5000 rpm, 4°C for 30 minutes. The expression pellets were resuspended in buffer A (20 mM Tris-HCl, pH 8.0, 100 mM NaCl, 10% glycerol, 5 mM β-mercaptoethanol). The cells were lysis by sonication; the lysate was centrifuged at 15000 x g for 30 minutes at 4 °C. The pellet thus obtained was resuspended in buffer B (1% Triton X-100, 20 mM Tris, pH 8.0) and again sonicated and centrifuged. The Pellet was further washed with buffer C (1 M NaCl, 20 mM Tris, pH 8.0 and centrifuged; the washed pellet was finally dissolved in Buffer D (8 M urea, 20 mM Tris, pH 8.0) overnight at room temperature. The lysate was centrifuged, and the supernatant was loaded onto a Nickel-NTA affinity column, which was pre-equilibrated with buffer D. For in-column refolding, protein bound to the Ni^2+^-NTA column was washed with buffer D by gradually decreasing the urea concentration from 8 M to 1 M with an interval of 1 M. At the end of the gradient, the column was washed with 20 ml of buffer A containing 20 mM imidazole. To remove impurities, the protein bounded column was washed with 10 ml of 20 mM Tris-HCl, pH 8.0, 100 mM NaCl, 5 mM β-mercaptoethanol, 30 mM imidazole, 5 % v/v glycerol. The protein was eluted in buffer A containing different gradients of imidazole (100 mM, 300 mM, 500 mM, 700 mM, 800 mM). The purity of the purified protein was analyzed by electrophoresis on a 15% polyacrylamide SDS gel. Fractions containing protein were pooled and were dialyzed with buffer containing 20 mM Tris-Cl, 100 mM NaCl, pH 8.0 (buffer E) to remove imidazole. The protein concentration was determined by measuring absorbance on Bio-Spectrometer and using the molar extinction coefficient of 19730 M^−1^ cm^−1^ at 280 nm.

The oligomeric status of the purified protein was determined by size exclusion chromatography using Superdex 200 Increase (10/300) GL column in a GE Akta purifier FPLC system. The molecular mass standards were dissolved in 20 mM Tris-HCl, pH 8.0, 100 mM NaCl, and were used to calibrate the column. To check the elution profile, WhiB6 (1.0 mg ml^-1^) was dialyzed against buffer E and was loaded to the pre-equilibrated column using a loop of 1 ml and injection volume of 800 μl. A flow rate of 0.5 ml min^-1^ was maintained.

### Preparation of apo WhiB6

The cysteine-bound iron-sulfur cluster [4Fe-4S] present in all the seven WhiB proteins (holo form) is oxidation labile; they disintegrate under oxidative conditions. However, the rate of degradation of the cluster varies among WhiB proteins, probably due to the difference in the extent to which the cluster is exposed to the external environment (23). Moreover, it has been reported that the binding efficiency of WhiB proteins (WhiB1, WhiB2, WhiB3, and WhiB4) to DNA is higher in their apo form than in the holo form (8, 11, 26, 27). Additionally, the apo form is the active form of WhiB proteins. Hence, to ensure that the iron-sulfur cluster has been completely removed, the purified protein was incubated with EDTA and potassium ferricyanide in a molar ratio of protein: EDTA: ferricyanide in 1:50:20 at 25°C. After 20–30 minutes or after the extensive loss of color, the solution was dialyzed in Buffer E for approximately 20 hours (28). The apo-WhiB6 protein thus obtained was further used in all the experimental studies.

### Insulin disulfide reductase assay

The reduction of insulin disulfides by DTT catalyzed using WhiB6 protein was analyzed using Holmgren’s (1979) method with some modifications (29). Apo WhiB6 protein of different concentrations (10 uM, 15 uM, 20 uM) was used to study the reductase assay. The stock solution of insulin was prepared in 20 % acetic acid. The stock solution concentration was determined using an extinction coefficient of 6000 M^-1^ cm^-1^ at 280 nm. The protein was pre-incubated for 1 hour in buffer E supplemented with 1 mM DTT, then 0.13 mM insulin was added to begin the reaction. To monitor the precipitation of the reduced insulin chains, absorbance at 600 nm every 30 seconds was recorded in a ThermoScientific Varioskan LUX Multimode Microplate Reader until A_600_ reached the saturation level. Insulin in the presence of DTT was used as a control for the uncatalyzed reduction of insulin by DTT. Buffer E supplemented with DTT was taken as blank.

### Biophysical characterization of apo-WhiB6 interaction with *espA* promoter DNA

For the WhiB6-*espA* promoter DNA interaction study, a 90 bp sequence of *espA* promoter DNA was used. Two strands of the promoter DNA were outsourced by GCC Biotech India Pvt. Ltd., West Bengal, India. The sequence of *espA* promoter DNA used was 5’-TGCCGTCTTCACGAACTCAAAACTGACGATCTGCTTAGCATGAAAAAAACTGTTGACATCGGCCAAGCATGACAGCCAGACTGTAGGCCC-3’ and 5’-ACGGCAGAAGTGCTTGAGTTTTGACTGCTAGACGAATCGTACTTTTTTTGACAACTGTAGCCGGTTCGTACTGTCGGTCTGACATCCGGG-3’. The single strands of the promoter DNA obtained were annealed to get double strands. Equal volumes of the equimolar oligonucleotides were mixed in an annealing buffer (10 mM Tris, pH 7.8, 50 mM NaCl and 1 mM EDTA). The reaction was incubated at 95 °C for 5 minutes, allowed to cool down gradually to room temperature, and cooled to 4 °C for temporary storage. The microtube was briefly centrifuged (short spun) to draw all the moisture away from the lid (30). For ITC experiments, the annealing buffer was 20 mM Tris, pH 8.0, 100 mM NaCl; to maintain similar buffer composition for the protein (WhiB6) and the ligand (*espA* promoter DNA).

The change in the secondary and tertiary structure of the WhiB6 on binding to nucleic acid was investigated by far-UV circular dichroism and steady-state intrinsic tryptophan fluorescence spectroscopy, respectively; the thermodynamic parameters were obtained using isothermal titration calorimetry (ITC); surface-enhanced Raman spectroscopy (SERS) gave insight into the conformational change and the residues involved in the interaction. Interaction studies was done under different conditions: Interaction between oxidized ((S)_2_) and reduced ((SH)_2_) WhiB6 and the promoter DNA was studied in the presence and absence of MgCl_2_ in the binding buffer.

### Steady-state Fluorescence spectroscopy

The binding affinity of WhiB6 with *espA* promoter DNA in the presence and absence of magnesium chloride (MgCl_2_) (10 mM) was determined from steady-state fluorescence quenching measurements. Fluorometric measurements were carried out on a Varian Cary Eclipse fluorimeter. The intrinsic tryptophan fluorescence emission spectra were recorded between 310 and 520 nm, keeping the excitation wavelength at 295 nm. The excitation and emission slit widths were both set at 5 nm. 15 μM Apo WhiB6 protein in 20 mM Tris HCl (pH 8.0), 100 mM NaCl buffer was incubated with increasing promoter DNA concentrations (0 μM, 1 μM, 2 μM, 3 μM, 4 μM, 5 μM, 6 μM, 7 μM, 8 μM, 9 μM and 10 μM) for 5 minutes.

The decrease in fluorescence intensity at ∼350 nm, after subtracting the contributions from the promoter DNA alone, was fitted using the modified Stern-Volmer equations to obtain the values of binding parameters like *n* (number of binding sites) and *K*_*b*_ (binding constant).

### Quenching data analysis

The quenching data obtained were analyzed by the modified Stern-Volmer equation; the binding affinity of the protein-DNA interaction and the number of binding sites were calculated (31). The equation is given by

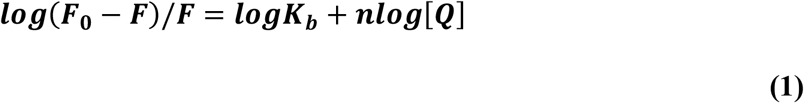

where *F*_0_ and *F* are the respective fluorescence intensities of fluorophore in the absence and presence of the quencher, [*Q*] is the quencher concentration, *n* is the number of binding sites for quencher on protein, and *K*_*b*_ is the binding constant. The slope and intercept of the plot between 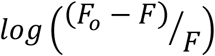 and *log*[*Q*] give the value of *n* and *K*_b,_ respectively.

The fluorescence data of WhiB6-espA interaction was corrected for the inner filter effect (IFE) of *espA* promoter DNA on the emission intensity of the protein using equation 2.

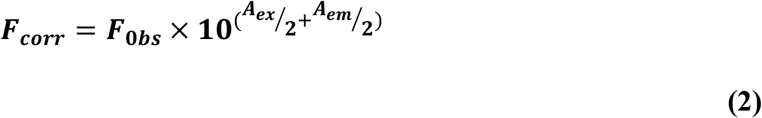

Where, F_corr_ is the IFE-corrected fluorescence emission intensity, and F_obs_ is the observed (uncorrected) fluorescence emission intensity. A_ex_ and A_em_ are the absorbance of the sample at excitation and emission wavelengths, respectively (32).

### Isothermal Titration Calorimetry (ITC)

ITC experiments were carried out using a MicroCal iTC200 titration microcalorimeter (Malvern) at 25 °C. All the samples were prepared in 20 mM Tris-Cl, pH 8.0, 100 mM NaCl (in the presence and absence of MgCl_2_ (10 mM). The samples were filtered and degassed before the measurements. 20 μM WhiB6 (in the cell) was titrated with 200 μM *espA* promoter ds-DNA in the syringe in a series of 20 injections of 2 μl each. Relevant reference experiments were performed by titrating the *espA* promoter DNA in the syringe into the buffer (with and without MgCl_2_). ITC data were analyzed using Origin 7.0 software and fitted with a one-binding site model.

### Far-UV Circular Dichroism (CD) spectroscopy

Far-UV CD measurements were performed at 25 °C using AVIV Model 420SF Circular Dichroism Spectrophotometer (Biomedical, inc. Lakewood, NJ USA) equipped with a peltier temperature controller system. The apo-WhiB6 protein was taken at a concentration of 15 μM in 100 mM NaCl, 20 mM Tris HCl (pH 8.0) buffer in the presence and absence of MgCl_2_. The appropriate volume of DNA stock was added to get a final DNA concentration of 1 μM, 2 μM, 3 μM, 4 μM, 5 μM, 6 μM, 7 μM, 8 μM, 9 μM, and 10 μM in the samples prepared for the CD. The incubation was done for 5 minutes. Three scans of far-UV CD were taken in a quartz cuvette of path length 1 mm, from 260 to 190 nm in increments of 1.0 nm with a scan rate of 20 nm/min and averaged. The baseline (buffer alone) and control spectra of MgCl_2_ and DNA alone were subtracted from the spectra of WhiB6-DNA-MgCl_2_ samples. The data were analyzed using the Origin 7.0 software.

The effect of WhiB6 protein on the conformation of *espA* promoter DNA was studied using Far-UV CD spectroscopy. The *espA* promoter DNA was taken at a fixed concentration of 4 μM in 20 mM Tris HCl, 100 mM NaCl, and pH 8.0 buffer in the presence of 10 mM MgCl_2_. The concentration of WhiB6 was varied as 0 μM, 4 μM, 8 μM, 12 μM, 15 μM, and 20 μM. The samples were incubated for 5 minutes. Three scans of far-UV CD were taken in a quartz cuvette of path length 1 mm, from 360 to 200 nm, in increments of 1.0 nm with a scan rate of 20 nm/min and averaged. The baseline (buffer alone) and control spectra of MgCl_2_ were subtracted from the spectra of WhiB6-DNA-MgCl_2_ samples. The data were analyzed using the Origin 7.0 software.

### Raman Spectroscopy and SERS

Plasmonic Silver (Ag) nanoparticles were prepared using the Lee-Meisel method (33). 18 mg of AgNO_3_ was added to 50 ml of Millipore-Q water with continuous stirring, brought to boiling, followed by the addition of 1% solution of sodium citrate while constant stirring. The nanoparticles were characterized by a Shimadzu UV-Visible spectrometer, transmission electron microscopy (TEM), and dynamic light scattering (DLS). The Raman and SERS spectra of the samples were recorded using a Horiba Xplora Raman spectrometer. Typically, the concentration of protein used in the experiment was in the micromolar range, and the molar concentration ratio of nanoparticles to the protein molecules was 1:100. 10 microliters of the solution of the complex was dropped on a quartz slide for acquiring the spectra. The excitation wavelength was set to 785 nm, and a 100x high NA objective was used to collect the spectra. The hole, slit width, and grooves for all samples were set to 300, 100, and 1200 lines/mm, respectively, and the typical accumulation time was 30 seconds. The post-processing of the raw spectra, including background subtraction and smoothening, was done using the Origin 7.0 software.

## RESULTS AND DISCUSSION

### WhiB6 protein was purified from inclusion bodies using the urea denaturation method

WhiB6 protein expressed in BL21(DE3) cells of *E. coli* at 37 ºC was expressed in the inclusion bodies. The urea-denaturation method was employed to purify it from the inclusion bodies. Due to the presence of a histidine tag (His_6_) at the C-terminal of the protein, Ni-NTA affinity chromatography was used for the purification. In-column refolding worked well for refolding the protein. On gradually decreasing the urea concentration, the protein was refolded inside the column. It was confirmed by the shift in wavelength maxima of the fluorescence spectra of the urea-denatured protein and that of the protein eluted from the column. The λ_max_ of the urea-denatured protein was observed at 354 nm, whereas the in-column refolded protein had its wavelength maxima at 350 nm (Figure 1A); the blue shift in the λ_max_ suggests the refolding of the protein. This observation was in accordance with the in-column refolding data reported by S. Alam et al. for WhiB4 (28). The purified WhiB6 protein was analyzed over SDS-PAGE; it corresponded to a molecular weight of ∼11 kDa (Figure 1B).

**Figure 1:**
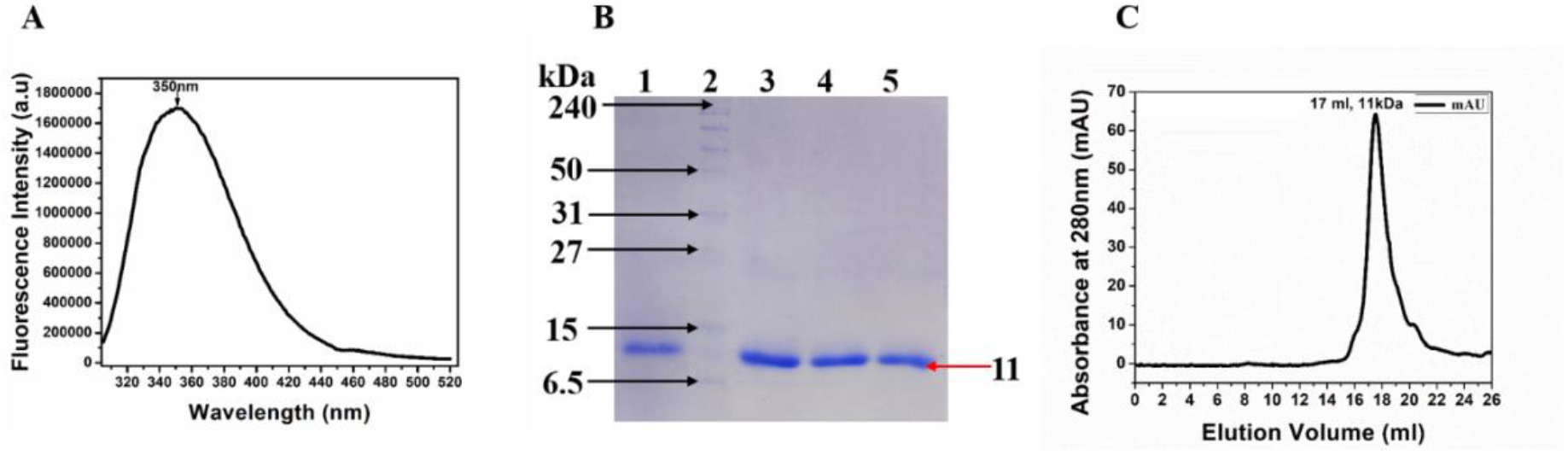
(**A**) Intrinsic Tryptophan fluorescence emission spectra of In-column refolded WhiB6. (**B)** 12% SDS-PAGE gel image showing purification profile of WhiB6 purified by Ni^2+^-NTA affinity column. Lane 1, 3, 4 & 5, fractions containing purified protein; lane 2, standard protein marker. (**C**) Gel filtration chromatogram of WhiB6 showing the elution profile of purified WhiB6 on Superdex 200 Increase (10/300) GL column; single peak shows the presence of monomer. The elution volume (17 ml) of protein corresponded to the size of ∼11 kDa.

The size exclusion profile of WhiB6 affirmed that the purified protein was in a monomeric state (∼11 kDa). Upon comparing the elution volume (17 ml) with the calibration curve plotted using the standard proteins (Supplementary Figure S1), the apparent molecular weight of the WhiB6 protein was determined. It corresponded to the molecular mass of a monomer (molecular weight of WhiB6 ∼11 kDa, Figure 1C).

### Insight into the Secondary structure of Apo-WhiB6 protein

The secondary structure of the apo-WhiB6 protein was studied using circular dichroism spectroscopy and Raman spectroscopy. The far-UV CD spectra revealed that the apo-WhiB6 has more of the random coil conformation. It shows higher molar ellipticity at 202 to 204 nm, indicating a significant proportion of random coils (Figure 2A). The β-strands content was also present; however, the α-helical content was minimum. Analysis of the amide I region acquired through Raman spectra of protein, as shown in Figure 2B, showed a similar result as CD spectroscopy. The Raman spectral peak of WhiB6 corresponded to an intense Raman peak around 1638 cm^-1^ and 1647 cm^-1,^ indicating that the secondary structure of protein mainly consists of a random coil or unordered structure (34). Similarly prominent Raman peak around 1661 cm^-1^ indicated the α-helical structure (35). The band around 1681 cm^-1^ indicated the random coil structure, whereas the small bands around 1628 cm^-1^ also signified the β-sheet structure (36). The modes related to the α-helix overlap with that of other secondary structures. Hence, it can be concluded that the protein predominantly consists of a random coil. The secondary structure data obtained of apo-WhiB6 is in accordance with the information reported by Kudhair et al. (2017) in their supplementary Figure 5a. They stated that the holo-WhiB1 had a significant amount of α-helical content; however, the apo-WhiB1, on removing the iron-sulfur cluster, lost almost all its helical content and attained random conformation. They further hypothesized that this conformation alteration could be responsible for modulating DNA binding in the apo conformation (5).

**Figure 2:**
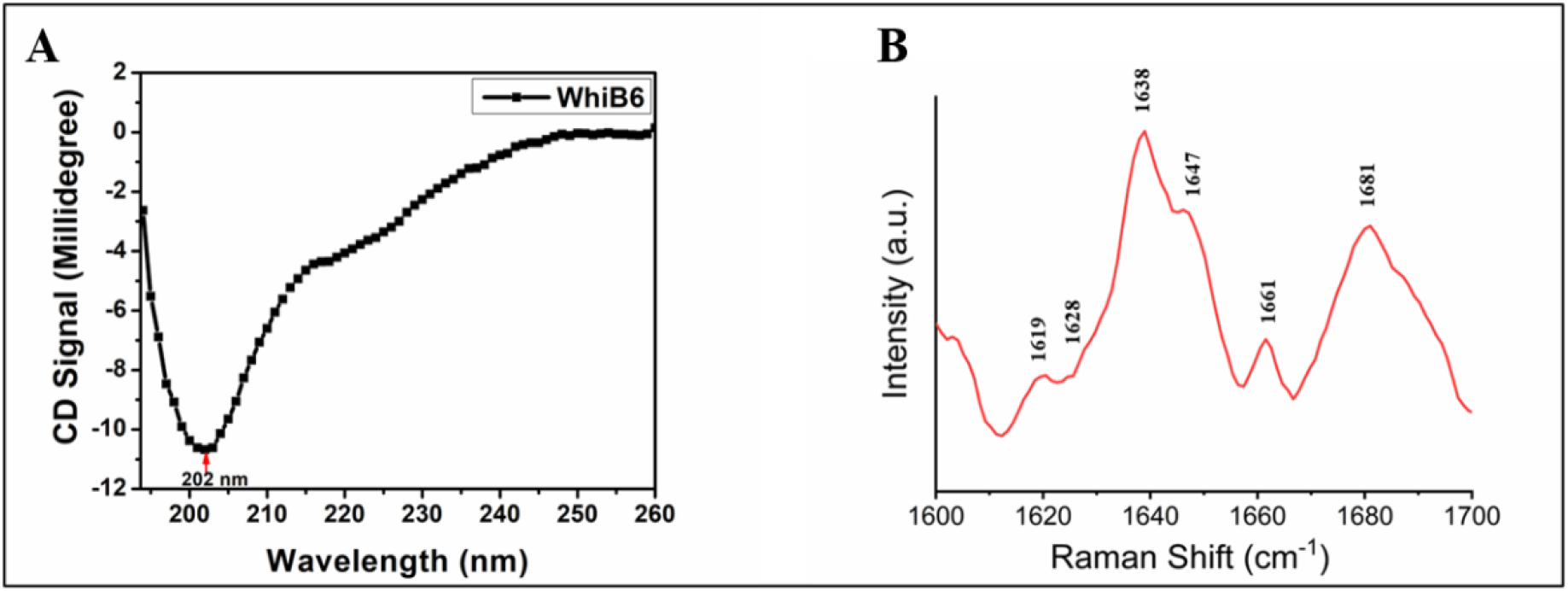
(**A)** Far-UV Circular Dichroism spectra of Apo-WhiB6. (**B**) Amide-I mode region of Raman spectra of WhiB6.

### WhiB6 catalyzed the reduction of the disulfide bond of insulin

Apo-WhiB proteins (except WhiB2) possess insulin disulfide reductase activity (23). The disulfide reductase activity of WhiB6 was investigated by the insulin disulfide reduction assay, where the test protein catalyzes the reduction of insulin by dithiothreitol DTT (29). Insulin comprises two polypeptide chains (A and B) interlinked by two disulfide bonds. The disulfide bond gets reduced in the presence of dithiothreitol (DTT). Upon reduction, the free B chain forms insoluble precipitates, leading to an increase in the absorbance at 600 nm. Thus, the onset and the rate of insulin aggregation are directly proportional to the protein’s reducing efficiency and concentration.

The reduction of insulin by DTT was determined in the presence and absence of apo WhiB6. On the addition of the WhiB6 to insulin + DTT reaction, the optical density at 600 nm increased compared to the uncatalyzed reaction (Figure 3). It showed that the purified apo-WhiB6 protein possessed insulin disulfide reductase catalytic activity. Furthermore, it was observed that the disulfide reductase activity of the WhiB6 protein was directly proportional to its concentration; as the concentration of the protein increased, its catalytic activity also increased (Figure 3).

**Figure 3:**
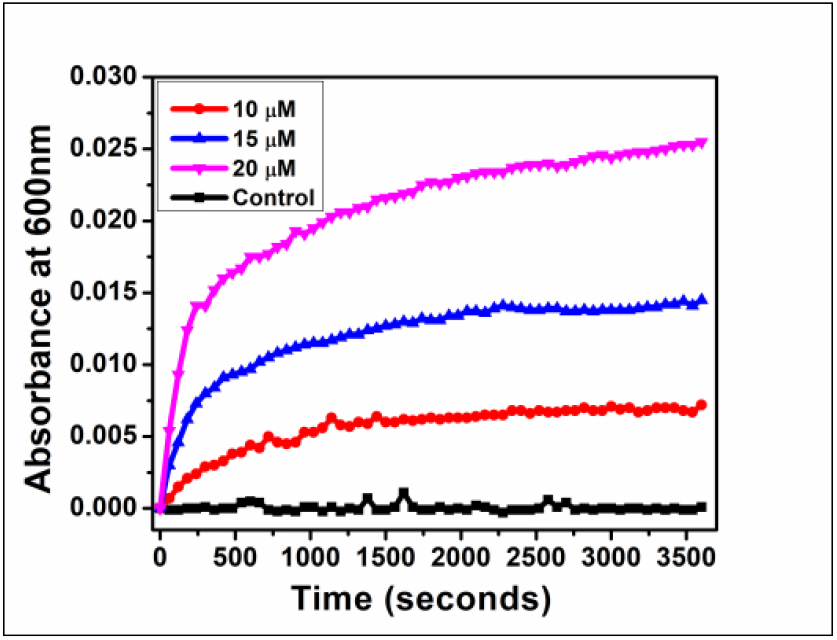
Insulin Disulphide Reductase assay of WhiB6.

### Investigating the interaction between WhiB6 protein and *espA* promoter DNA

The interaction between the WhiB6 and *espA* promoter DNA has been studied by circular dichroism spectroscopy (CD), Isothermal titration calorimetry (ITC), steady-state fluorescence spectroscopy, and Raman Spectroscopy. The interaction was studied under different conditions. WhiB proteins are redox-sensitive; therefore, the interaction was studied with both oxidized and reduced apo-WhiB6 proteins to develop an overall understanding. WhiB6 protein was incubated with 1 mM TCEP at 25 °C for 1 hour for reduction. Various salts with optimised concentrations are known to act as a driving force in forming stable protein-DNA complexes (37). Moreover, it is established that magnesium chloride (MgCl_2_), when added to the binding buffer, enhances the probability of protein-DNA interaction (8). It is also known to stabilize the structure of a protein and improve its affinity to DNA (38). For the same reason, 10 mM MgCl_2_ was used in the binding buffer for interaction studies. According to the previous reports, WhiB proteins (WhiB1, WhiB2, WhiB3, and WhiB4) bind to DNA with greater affinity in their apo form than the holo form (8, 11, 26, 27); therefore, all the interactions studies were carried out utilizing apo-WhiB6 protein.

### Steady-state fluorescence spectroscopy gave insight into the binding affinity of WhiB6 protein to *espA* promoter DNA

The structural changes in protein upon protein-DNA interaction can be determined by comparing the protein-DNA interaction spectra with the protein alone spectra (32). In the present study, the binding of the oxidized ((S)_2_) and reduced ((SH)_2_) WhiB6 protein with the *espA* promoter DNA in the presence and absence of MgCl_2_ in the binding buffer was studied using steady-state fluorescence spectroscopy. WhiB6 was titrated with an increasing concentration of *espA* Promoter DNA, and tryptophan fluorescence of the resulting solutions was recorded. The emission spectrum of WhiB6 in the absence of the promoter DNA showed an emission maximum at 350 nm when excited at 295 nm (in both oxidized and reduced conditions and in the presence and absence of MgCl_2_), which inferred that the Trp residue is exposed to the polar environment. The excitation wavelength of 295 nm was chosen to excite tryptophan and avoid any contribution from Tyr residue. Quenching of the fluorescence intensity (Figures 4A, 5A, 6A, and 7A) and a blue shift in the λ_max_ of the fluorescence emission (Supplementary Figure S2, S3, & S4) was observed with a gradual increase in the concentration of *espA* promoter DNA (from 0-8 or 10 μM) under all the binding conditions (oxidized and reduced WhiB6-*espA* promoter DNA interaction in the presence and absence of MgCl_2_) except during the reduced WhiB6-*espA* promoter DNA interaction in the absence of MgCl_2_ (a blue shift in the λ_max_ of the fluorescence emission was not observed) (data not shown). It was inferred that the environment and the polarity around tryptophan altered upon interaction. A similar result was obtained from SERS; protein-DNA interaction altered the 1364 cm^-1^ and 1387 cm^-1^ peaks corresponding to the tryptophan residues (Table 3) (Figure 11). This suggests that the interaction between the DNA and WhiB6 caused a significant change in the polarity and environment around tryptophan and in the protein’s tertiary structure.

**Figure 4:**
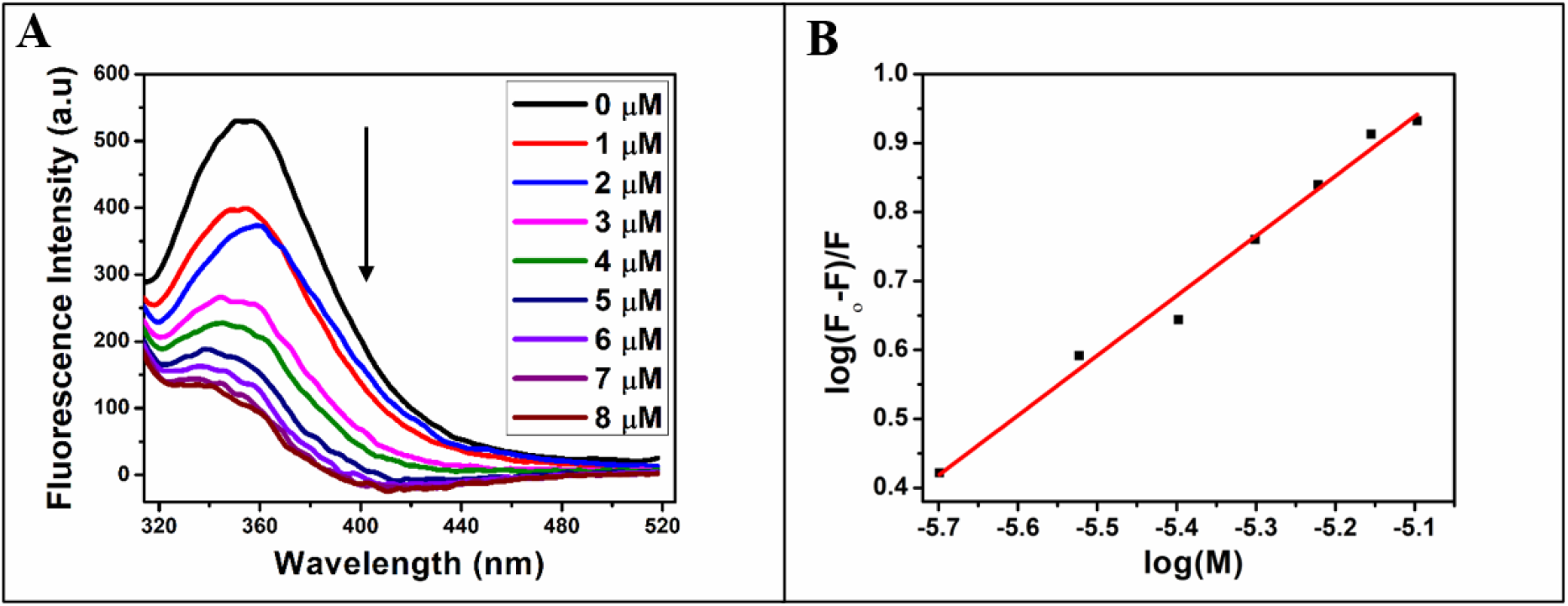
(**A**) Tryptophan spectra of WhiB6 (15 μM, pH 8.0) in the presence of *espA* Promoter DNA (in the **absence of MgCl**_**2**_ in binding buffer). **(B)** Modified Stern-Volmer plot to estimate the binding constant and number of binding sites. The emission maximum was obtained at 350 nm.

**Figure 5:**
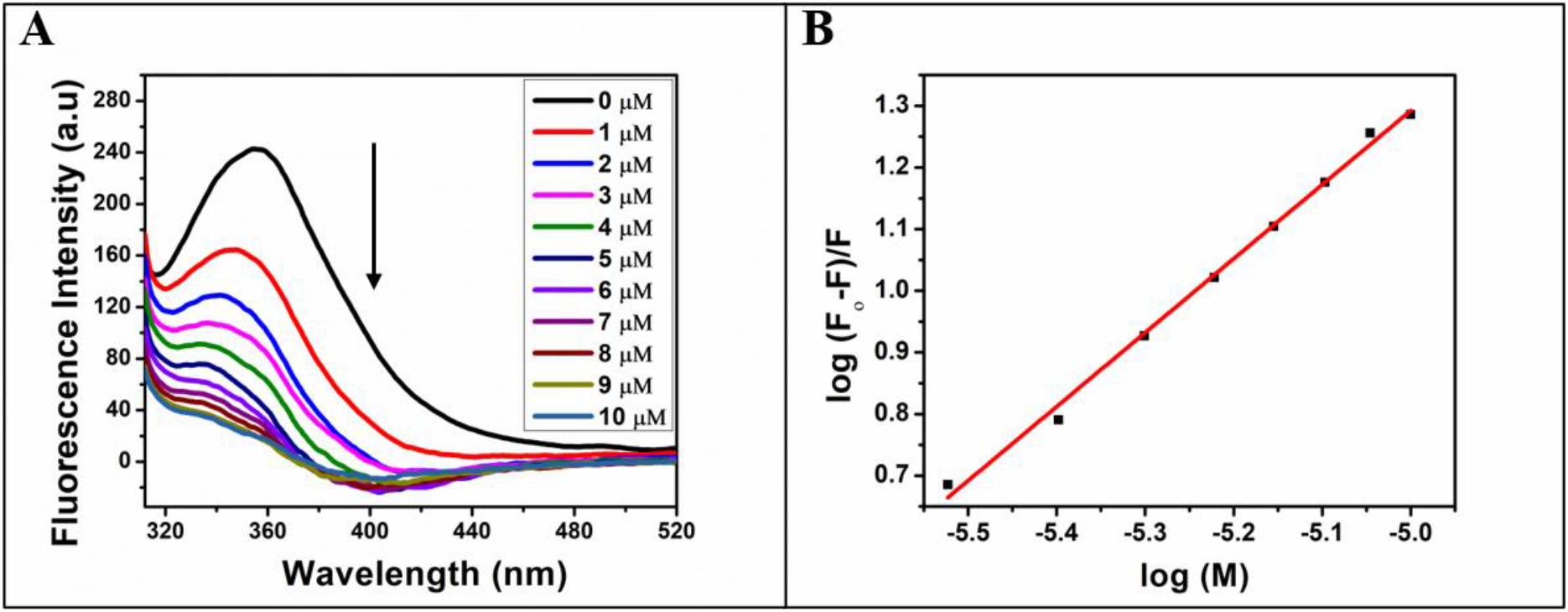
(**A**) Tryptophan emission spectra of WhiB6 (15 μM, pH 8.0) in the presence of espA Promoter DNA (in the **presence of 10 mM MgCl**_**2**_ in binding buffer) (**B**) Modified Stern-Volmer plot for the estimation of the binding constant and number of binding sites. The fluorescence emission maxima was observed at 350 nm.

**Figure 6:**
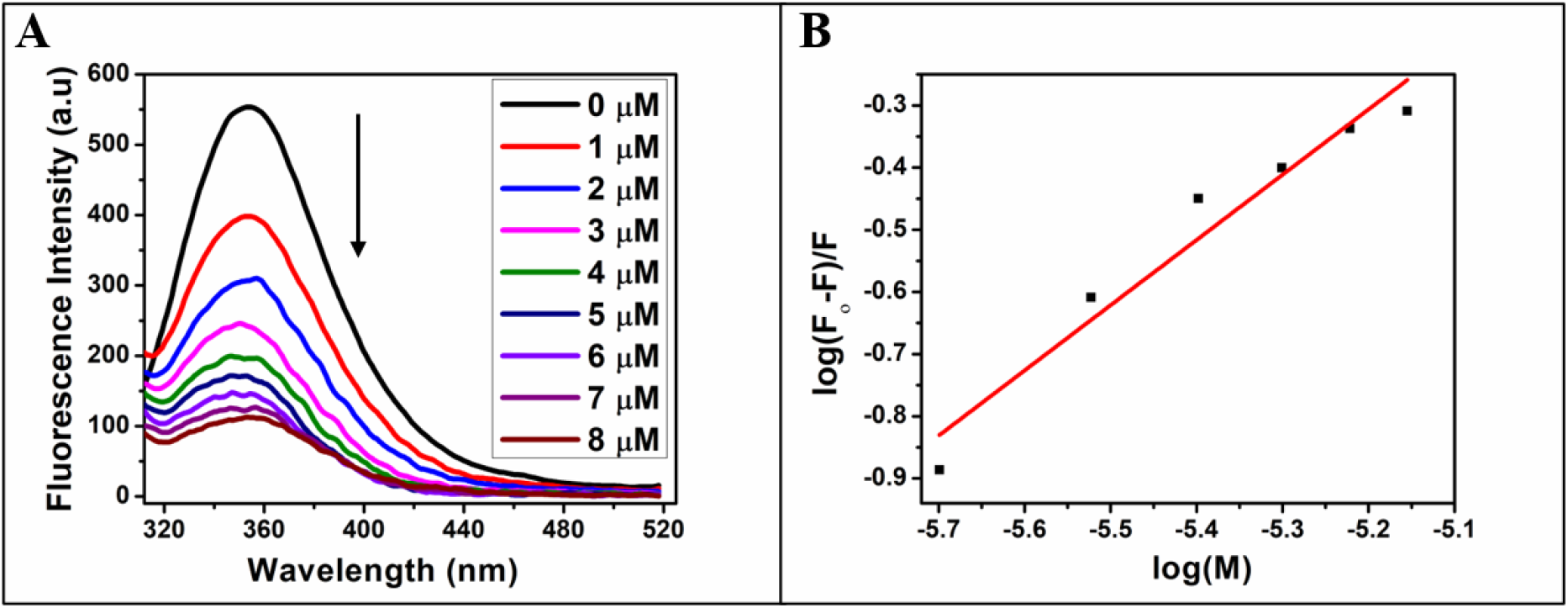
(**A**) Tryptophan spectra of **Reduced** WhiB6 (15 μM, pH 8.0) in the presence of *espA* Promoter DNA (in the **absence of MgCl**_**2**_ **in binding buffer**) (**B**) Modified Stern-Volmer plot to estimate the binding constant and number of binding sites. The fluorescence emission maxima was observed at 350 nm.

**Figure 7:**
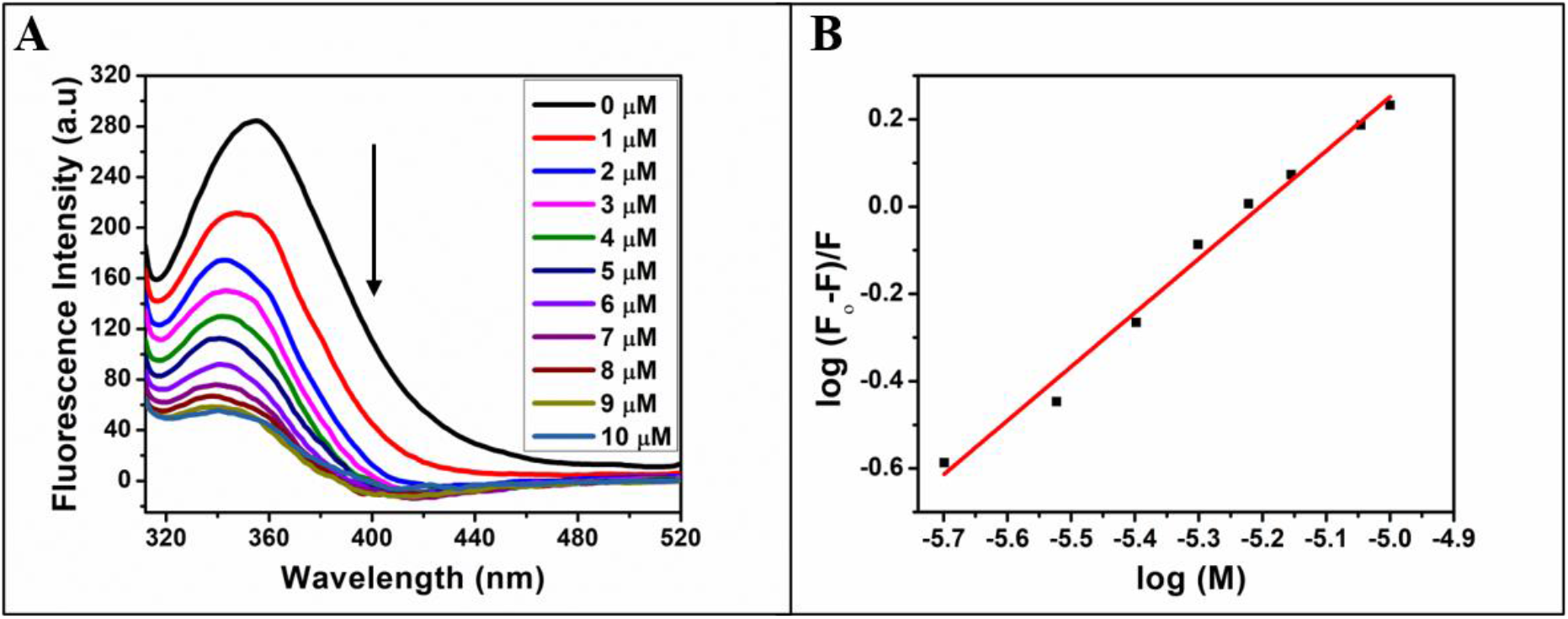
(**A**) Tryptophan spectra of **Reduced** WhiB6 (15 μM, pH 8.0) in the presence of *espA* Promoter DNA (in the **presence of 10 mM MgCl**_**2**_ in binding buffer) (**B**) Modified Stern Volmer plot of WhiB6 (15 μM, pH 8.0) for the estimation of the binding constant and number of binding sites at 25 °C. The fluorescence emission maxima was observed at 350 nm.

The inner filter effect (IFE) correction of the protein emission intensity was done by measuring the absorbance at the excitation and emission wavelength for each concentration of the DNA. IFE occurs when the ligand molecule used in the protein-ligand interaction studies absorbs light at the excitation and emission wavelength of the protein (32, 39). The corrected fluorescence emission intensity, F_corr_, was obtained using Equation 2 and was used to generate the modified Stern-Volmer plot for estimating the binding constant.

The binding constant (*K*_*b*_) was obtained using the modified Stern-Volmer equation (Equation (1), Table 1, Figures 4B, 5B, 6B, and 7B). The binding constant, *K*_*b*,_ in the oxidized ((S)_2_) condition was 1.97×10^7^ M^-1^ and 2.31×10^5^ M^-1^ in the presence and absence of MgCl_2_, respectively, in the binding buffer. Whereas, in the reduced WhiB6 condition, the *K*_*b*_ was 2.71×10^6^ M^-1^ and 1.42×10^5^ M^-1^ in the presence and absence of MgCl_2_, respectively (Table 1).

**Table 1:** Parameters obtained by fitting of modified Stern-Volmer plot (in the **presence and absence of 10 mM MgCl**_**2**_ in binding buffer).

|  |  | Modified Stern-Volmer method for number of binding sites and binding constant |  |
| --- | --- | --- | --- |
| | | Binding constant, $K_b$ ($M^{-1}$) | Binding site value, $n$ |
| Oxidized WhiB6 | Absence of $MgCl_2$ | $2.31 \times 10^5$ | 0.9 |
| | Presence of $MgCl_2$ | $1.97 \times 10^7$ | 1.05 |
| Reduced WhiB6 | Absence of $MgCl_2$ | $1.42 \times 10^5$ | 1.0 |
| | Presence of $MgCl_2$ | $2.71 \times 10^6$ | 1.1 |

The steady-state fluorescence results inferred that the association constant obtained at all conditions, $K_b$ , was $\sim 10^5$ to $10^7 M^{-1}$ , illustrating a moderate binding affinity between WhiB6 and *espA* promoter DNA (40). The stoichiometry of binding, $n$ , was nearly 1:1.

It was concluded from the steady-state fluorescence spectroscopy observation that the presence of MgCl_2_ in the binding buffer enhanced the binding affinity of WhiB6 to the promoter DNA. Moreover, the binding efficiency was better in the case of oxidized WhiB6 ((S)_2_) as compared to reduced WhiB6 ((SH)_2_). This observation was in accordance with the reported data, which stated that in the apo-oxidized state, WhiB proteins (WhiB1, WhiB2, WhiB3, and WhiB4) bind to DNA with higher affinity than the reduced state (8, 11, 26, 27). Similarly, in the case of WhiB6, the binding affinity was 10^7^ in the oxidized ((S)_2_) form, whereas it was 10^6^ in the reduced state in the presence of MgCl_2_.

### Thermodynamic profile of WhiB6-*espA* promoter DNA interaction

Isothermal titration calorimetry (ITC) is the most reliable and efficient technique to measure the thermodynamic parameters of binding such as binding constant/association constant (*K*_*a*_), stoichiometry (*n*), entropy change (ΔS), and enthalpy change (ΔH) (41).

The ITC data analysis revealed that the interaction between 20 μM WhiB6 and 200 μM *espA* promoter DNA in the presence of 10 mM MgCl_2_ was exothermic (Figure 8), with an association constant of 1.4 ×10^6^ M^-1^ at 298 K (Table 2). The thermodynamic profile of the interaction (large negative value of enthalpy change and small negative value of entropy change) showed that the binding was enthalpically favored. It further suggested that the electrostatic interaction played a crucial role in stabilizing the complex. The negative ΔG indicated that the binding process was a spontaneous reaction. The stoichiometry of binding (*n*) was ∼ 1:1, suggesting one WhiB6 bind with one *espA* promoter DNA. The binding constant was 10^6^ M^-1^, indicating a moderate binding affinity between WhiB6 and the DNA (40), which is in accordance with the steady-state fluorescence interaction result. The ITC experiment was also conducted in the absence of MgCl_2_ in the binding buffer and in the reduced conditions; however, the curve obtained could not be fitted with any binding model. It was further inferred from the ITC data that the presence of MgCl_2_ facilitated better interaction between WhiB6 and *espA* promoter DNA.

**Figure 8:**
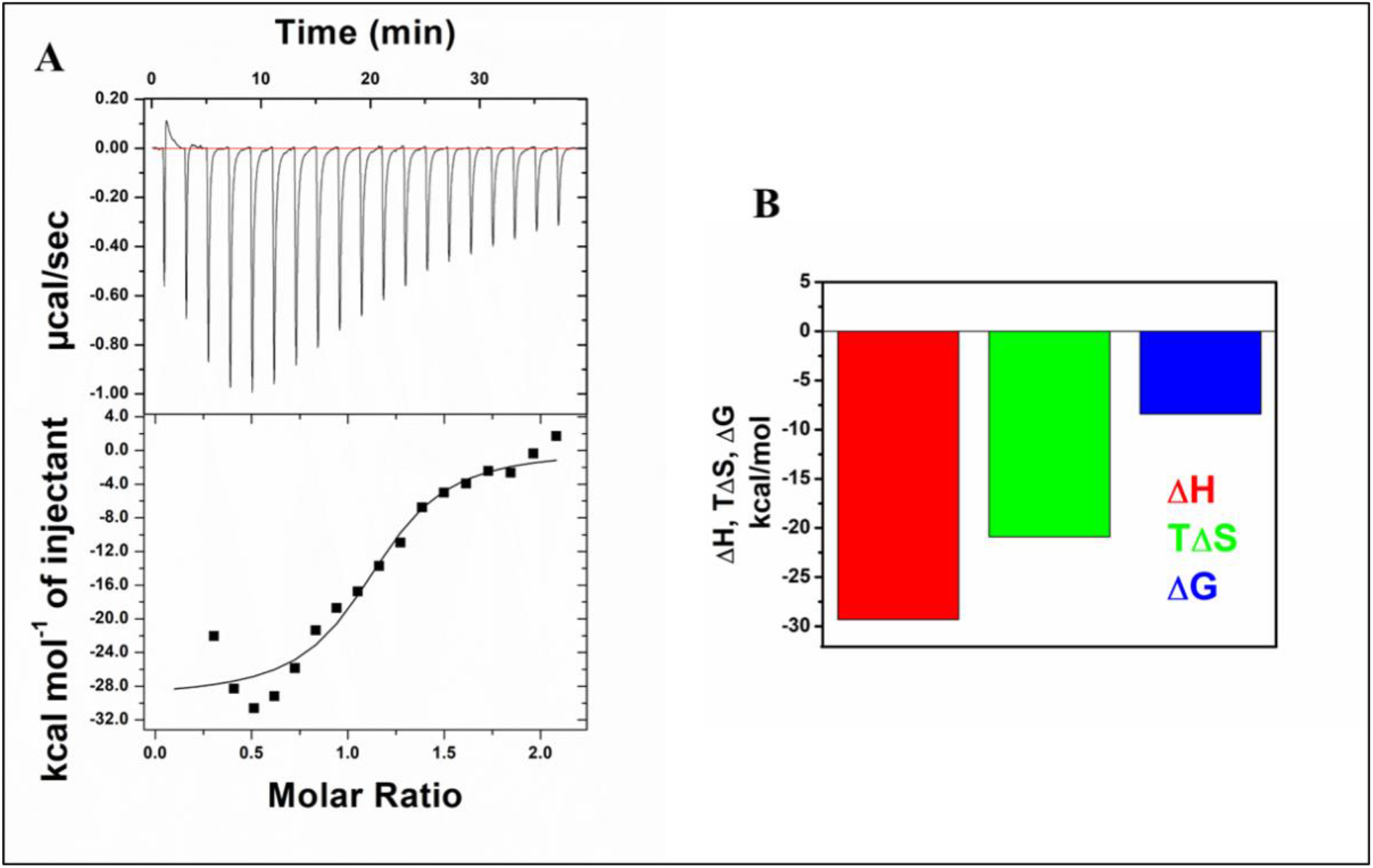
(**A**) Isothermal titration calorimetry (ITC) of interaction between WhiB6 (20 μM) and *espA* promoter DNA (200 μM) in 20 mM Tris pH 8.0, 100 mM NaCl, 10 mM MgCl_2_. The upper panel represents the raw ITC thermogram for heat flow during the reaction versus time, whereas the lower panel represents the integrated and normalized heat of *espA* promoter DNA titration into the WhiB6 solution. The solid line represents the best fit of the integrated data to a single binding site model. (**B**) Bar diagram comparing thermodynamic parameters in the binding events (in the presence of 10 mM MgCl_2_).

**Table 2:** The thermodynamic parameters for the interaction between WhiB6 and *espA* promoter DNA obtained from the isothermal titration calorimetric measurements performed at pH 8.0, 25 °C in the presence of MgCl_2_.

|  | N | K<sub>a</sub> (M<sup>-1</sup>) | ΔH (cal/mole) | ΔS (cal/mole/°C) | ΔG (kcal/mole) |
| --- | --- | --- | --- | --- | --- |
| **Presence of MgCl<sub>2</sub>** | **1.1 ± 0.04** | **(1.4 ± 0.06) × 10<sup>6</sup>** | **-(0.29 ± 0.01) × 10<sup>5</sup>** | **-70.1 ± 0.0** | **-8.40** |

**Table 3:** SERS band assignment for protein and protein-DNA complex with AgNPs. The appearance of the new band is indicated by *, s-sharp, sh-shoulder, br-broad, w-weak.

| Protein (cm<sup>-1</sup>) | Protein-DNA complex (cm<sup>-1</sup>) | Proposed band assignment |
| --- | --- | --- |
| 724 | 731 | $\delta$ (COO<sup>-</sup>)/ Adenine Ring breathing |
| - | 804 | Tyr or $\nu_{as}$(C-S-C) (44) |
| 834 | 834(sh) | Tyr (45) |
| 860(s) | 860(very weak) | Tyr |
| 917 | 914 | $\nu$ (C-COO<sup>-</sup>)/Ribose C-C stretching ((45) or C-C stretch of proline ring or arginine (46–48) |
| - | 941* | $\nu$(C-COO<sup>-</sup>)/ $\alpha$ helical C-C stretching vibration or proline (49) |
| 1003 | - | Phenylalanine or tryptophan(trp) (46, 50, 51) |
| 1030 | 1031 | Phenylalanine $\nu_{18a}$/C-N stretch |
| 1130(s) | 1123(br) | $\nu_{as}$ (CαCN) or C-H def or C-N stretching/Trp |
| 1185 | 1173 | $\nu_{as}$ (CCN), Tyr or Phe<sub>9a</sub> or Thymine (44, 52) |
| 1216,1245 | 1245 | His or amide III of β-sheet (52) or Trp |
| 1301 | 1296 | $\nu_{as}$ COO<sup>-</sup> or CH deformation or amide III-helix |
| 1334(weak) | 1334(sh) | Adenine or guanine ring modes (45) or Trp, (Cα-H<sub>def</sub> of α-helix (53) or $\delta$ (CH) (54) |
|  | 1364,1387 | Trp, COO<sup>-</sup> stretching |
| 1603, 1622 | 1616(s) | Phe or Tyr aromatic vibration (52) |
| 1658 | - | Amide I - α-helix |
| 1680 | 1678(br) | Amide I – random coil |

The slight difference in the binding parameters obtained from ITC and those obtained from the steady-state fluorescence measurements could be due to the difference in the working principles of both techniques. Steady-state fluorescence measures the local changes in the environment of fluorophores, whereas ITC measures the overall change in heat accompanying the binding process.

### Structural modulation of Secondary structure of WhiB6 upon interaction with *espA* promoter DNA

CD spectroscopy was used to investigate the alteration in protein structure on the interaction with DNA or vice versa. The asymmetric carbon atoms present in the sugars of nucleotides and all the amino acids (except for glycine) result in nucleic acids and proteins displaying optical activity (42). Nucleic acids dominate CD spectra between 250 and 300 nm. The region below 250 nm provides important information regarding the secondary structures of proteins complex with nucleic acids (42); hence the far-UV CD spectra for the WhiB6-DNA interaction study were recorded from 190 nm to 260 nm. A decrease in the CD signal was observed with an increase in the DNA concentration at all the conditions of the binding buffer, indicating small changes in the overall secondary conformation of WhiB6 protein on interaction with *espA* promoter DNA in the presence (Figure 9) and absence (Supplementary Figure S5) of MgCl_2_.

**Figure 9:**
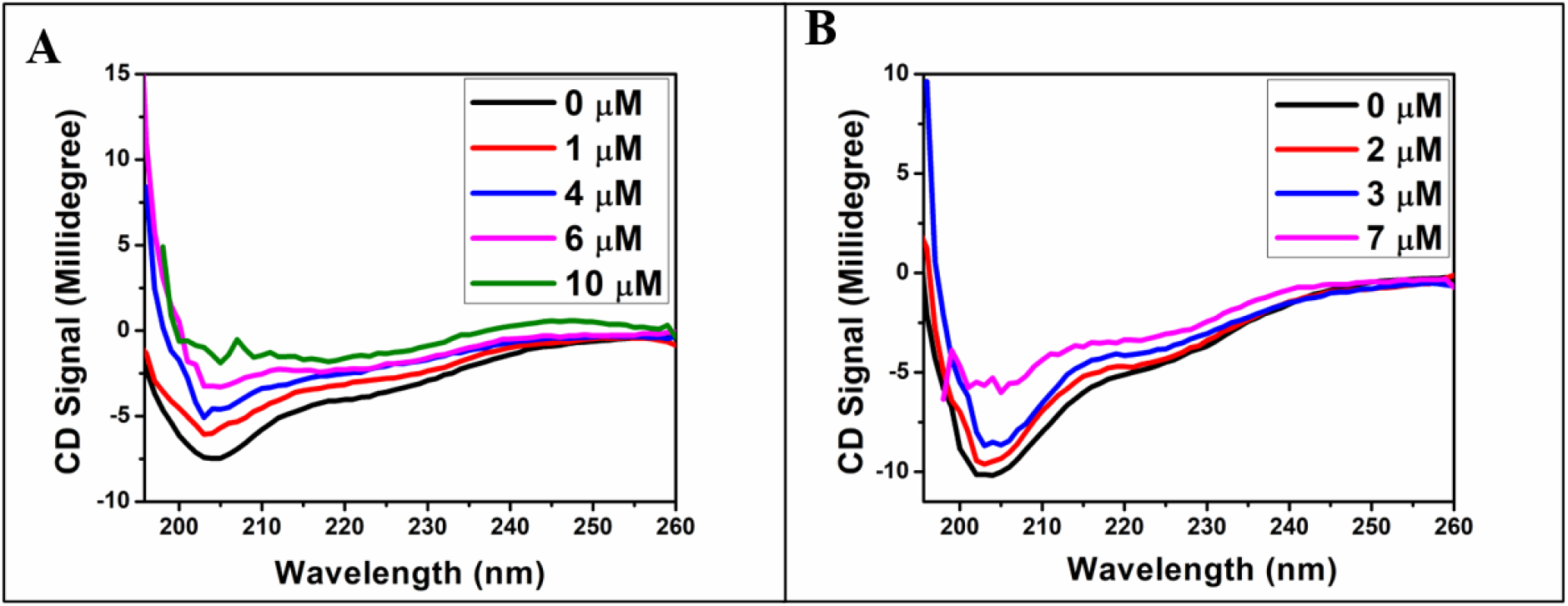
Far-UV Circular Dichroism spectra showing changes in the secondary structure of WhiB6 on interacting with espA promoter DNA in the **presence of 10 mM MgCl**_**2**_. (**A**) WhiB6 interaction with espA promoter DNA. (**B**) Reduced WhiB6 interaction with espA promoter DNA.

CD spectra of DNA is typical of B-DNA with a negative and positive peak at 250 and 280 nm, respectively. The titration of *espA* promoter DNA (4 μM) with WhiB6 of different concentrations (0 μM, 4 μM, 8 μM, 12 μM, 15 μM, and 20 μM) resulted in no significant change in the far-UV CD spectra of the *espA* promoter DNA (Figure 10). It was inferred from the data that the secondary structure of the *espA* promoter DNA remained unaltered upon interaction with the WhiB6 protein.

**Figure 10:**
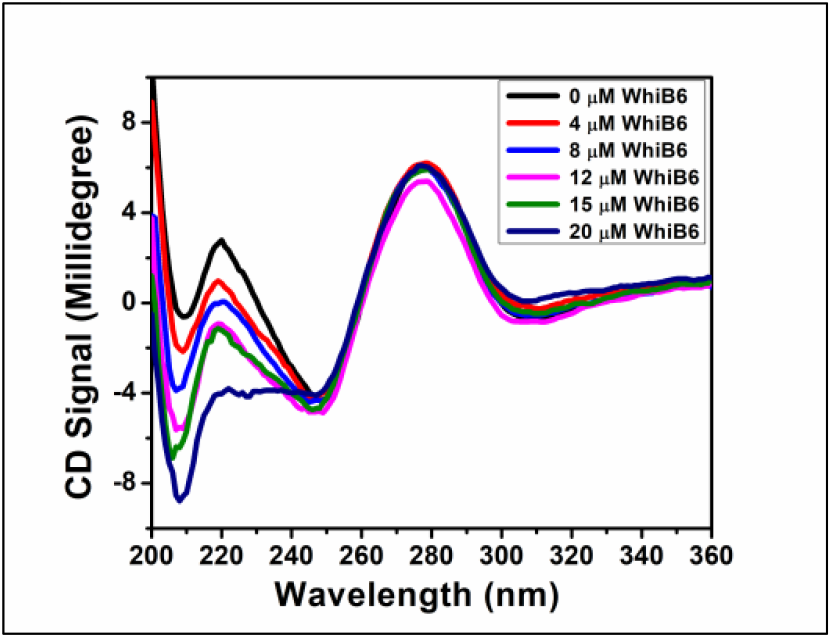
Far-UV CD spectra of *espA* promoter DNA titration with different concentrations of WhiB6 in the presence of 10 mM MgCl_2_.

A correlation was observed between the steady-state fluorescence spectroscopy, circular dichroism spectroscopy, and ITC result of the WhiB6-*espA* promoter DNA interaction. The fluorescence spectroscopy observation showed that until 4 μM DNA concentration, conformational change, as well as quenching, took place, as there was a significant blue shift in the maximum wavelength, λ_max,_ and a decrease in the fluorescence intensity. However, after 4 μM, there was no further blue shift in the spectra, and only fluorescence intensity decreased; It infers that there was no further change in the conformation; however, only quenching occurred due to binding. A similar change was observed in the CD spectra till 4-5 μM DNA concentration, conformational change was observed as the CD signal decreased, beyond 5 μM, the protein structure got distorted (Figure 9).

### SERS employed to study WhiB6-*espA* promoter DNA interaction

Surface-enhanced Raman spectroscopy (SERS) was used to analyze the interaction of WhiB6 with *espA* promoter DNA. SERS provides an alternative way to analyze complex biomolecules such as proteins in addition to various analytical techniques like infrared spectroscopy (IR), Circular dichroism (CD), X-ray crystallography, and Fourier transform infrared spectroscopy (FTIR), *etc*. (43, 44). Particularly, SERS is useful for probing proteins and determining the secondary structure conformations and interaction with ligand molecules, metal ions, and DNA in a non-destructive way involving minimal sample preparation (45).

The SERS spectra are dominated by residues that are in close proximity to the nanoparticle surface as the electromagnetic field intensity is a function of distance from the surface and decays down exponentially, thus giving unique surface information. Any binding events eventually lead to a change in the SERS spectra due to the change in orientation due to the binding or the modes of the ligand itself (36).

In the SERS spectra shown in the protein, DNA, and protein-DNA complex, a change in modes in terms of intensity and position was observed due to the binding of DNA with protein. Due to the enhancement of a few modes preferentially in the complex and the disappearance of some modes in the spectra of the complex, we inferred that a binding event is indeed taking place (Figure 11A). We observed that the binding of DNA to protein resulted in a decrease in the intensity of the amide I region corresponding to α-helix, but the β-sheets modes were more enhanced. The amide I band corresponding to α-helix and β-sheets lies in the regions of 1655-1662 cm^-1^ and 1670-1674 cm^-1^, respectively (56). The amide I region corresponding to the random coil is around 1680-1685 cm^-1^ (57). The band around 914 cm^-1^ can be attributed to proline or arginine ring vibrations or ribose C-C stretching and was seen to be altered in the spectra of the protein-DNA complex. The binding of DNA also resulted in the changes in intensities of modes related to tyrosine (Tyr) around 860 cm^-1^ and the emergence of peaks around 916 and 942 cm^-1,^ which could be attributed to arginine (Arg) and proline (Pro). The protein-DNA complex formation also results in changes in intensities in the region of 1340-1360 cm^-1^ which can be attributed to tryptophan (Trp), histidine (His), or valine (Val). The phenylalanine (Phe) mode at 1003 cm^-1^ was also found to diminish in intensity on the interaction of DNA. Thus, it was inferred that the DNA binding site was not far from the nanoparticle binding site (both before and after DNA binding) due to the observance of a few modes of the DNA even in the protein-DNA complex. The change in spectral intensity of Trp and also residues such as Arg, Pro and Phe in complex indicates the role of these residues during the interaction with DNA. In the protein-DNA complex, there was a change in the relative intensities of various secondary structures which reiterates the complex formation (Figure 12, B and C).

**Figure 11:**
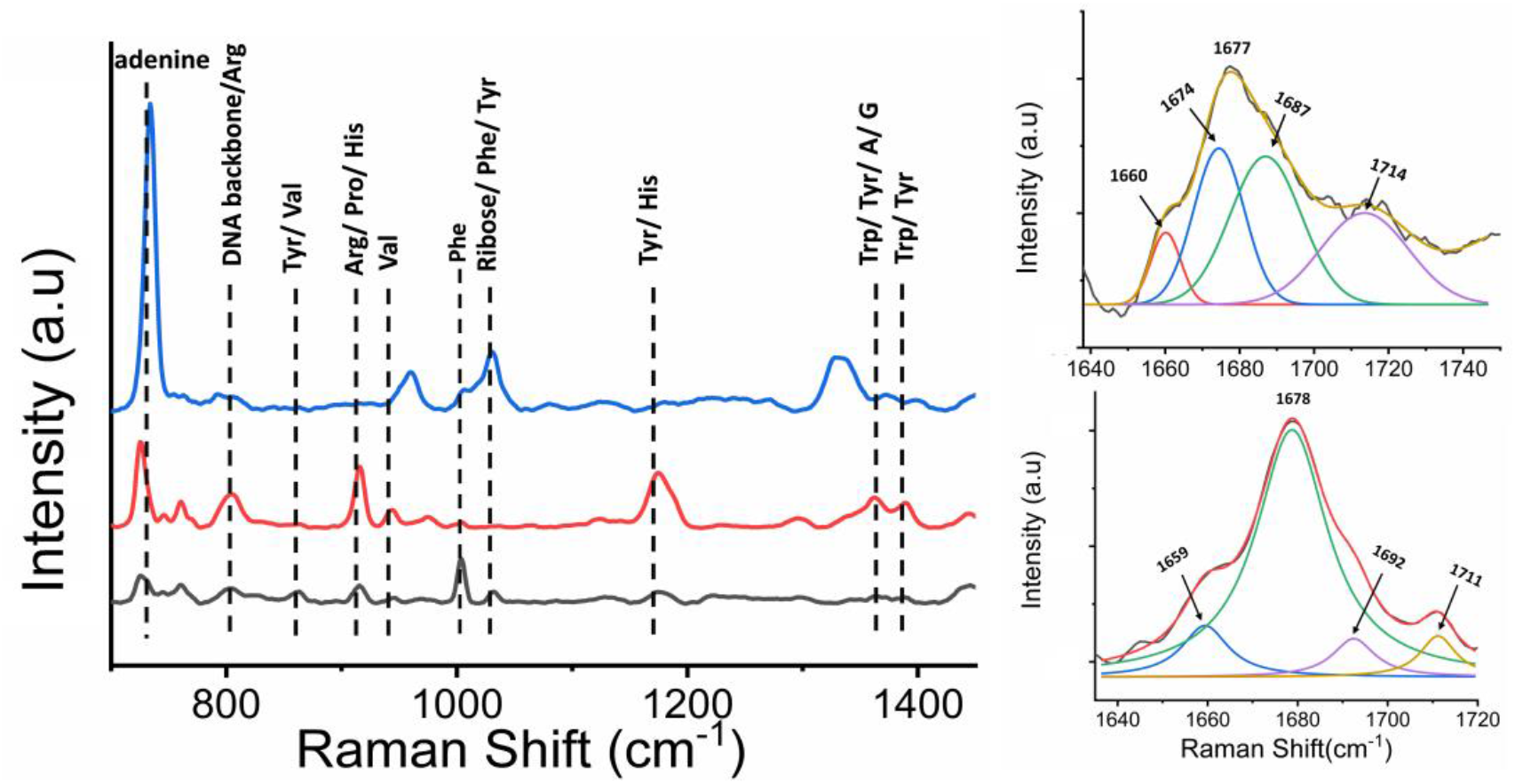
(**A**) SERS spectra of DNA (blue), complex (red), and protein (black). (**B**) Deconvolution of amide I band of protein-DNA complex. (**C**) Deconvolution of amide I band of protein.

**Figure 12:**
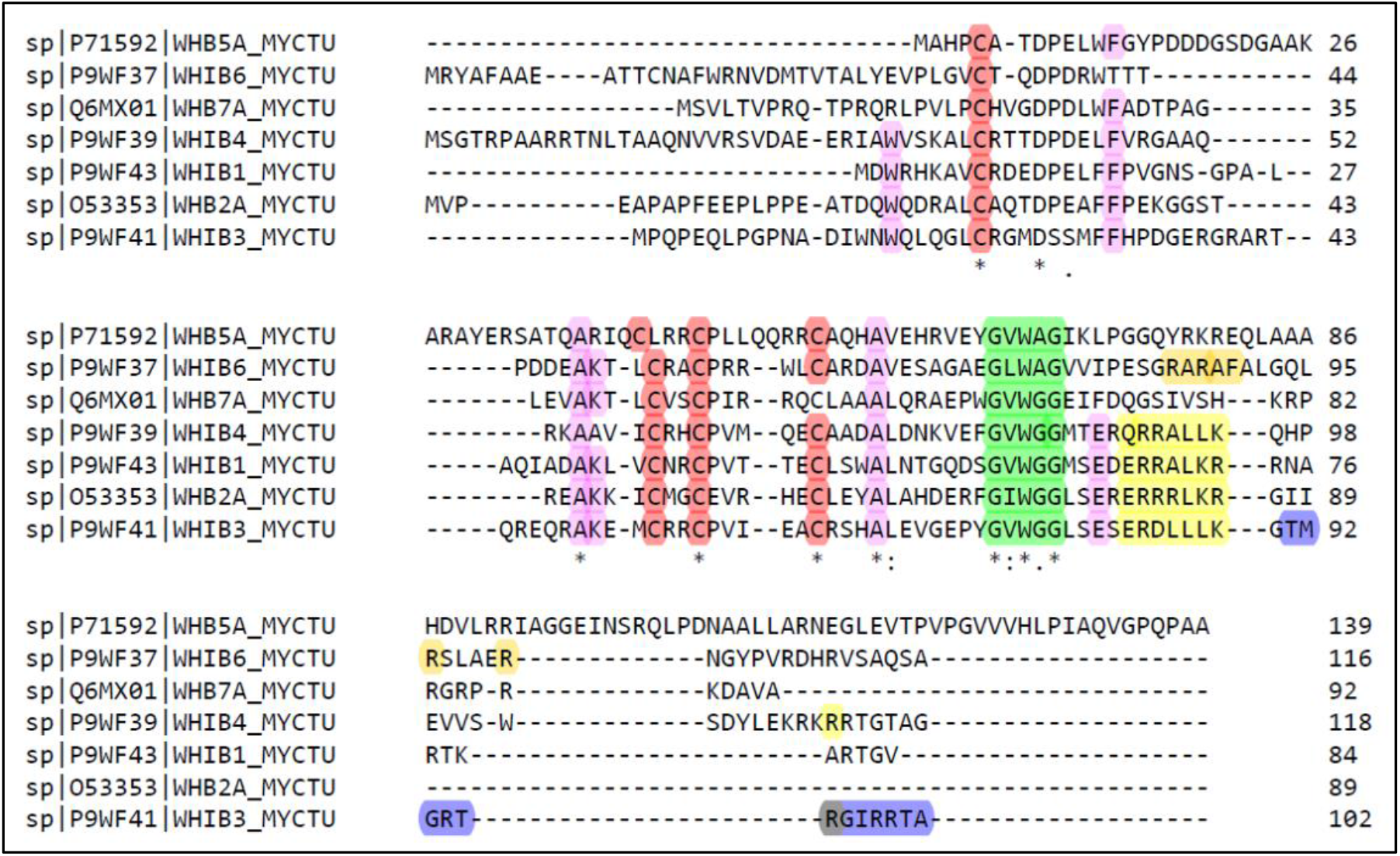
Multiple alignment results showing conserved WhiB6 sequence and regions. The conserved motifs have been highlighted; the four conserved cysteine residues are in red, the tryptophan region (G[V/I/L]W[G/A]G turn) in green, and the C-terminal basic amino acids in yellow, and other conserved residues are in pink.

The multiple sequence alignment result showed that WhiB6 is composed of the conserved domains (like other WhiB proteins): four conserved cysteine residues at the N-terminal domain, a (G[V/I/L]W[G/A]G motif or tryptophan region or β-turn and C-terminal domain consisting of basic amino acids that form helix-turn-helix (HTH) motif (Figure 12).

Zhai et al. in 2021 reported the solution structure of reduced apo-WhiB4. They showed that the reduced apo-WhiB4 had a disordered structure. The N-terminal domain was unstructured, but the C-terminal domain, responsible for DNA binding, had a preserved helix-turn-helix structure (58). The reduced apo-WhiB4 structural details were in accordance with the apo-WhiB1 NMR structure reported by Kudhair et al. in 2017. They also reported that the apo-WhiB1 lost most of the helical structure (compared to the holo form) and gained random coil conformation. However, the C-terminal helix retained the native-like folded conformation (5). According to the literature, the conformational change occurring upon conversion from holo to apo form (loss of the cluster) modulates the C-terminal helix to allow DNA binding activity (5).

Therefore, upon conversion to the apo form, WhiB proteins form an unstructured N-terminal domain, and a C-terminal domain consists of two α-helices forming an HTH motif. The four conserved cysteine residues are located in the N-terminal unstructured domain. The C-terminal HTH domain of the apo protein plays a crucial role in binding with DNA.

As mentioned above, the basic amino acids present at the C-terminal of the WhiB proteins form the helix-turn-helix like structure that is involved in binding with DNA (4, 27, 58). The residues predicted by SERS are also among the amino acids located at the C-terminal of the WhiB6 protein sequence; furthermore, some of them are positively charged, like arginine (highlighted in the protein sequence in Figure 13 in red color). Hence it further suggests that the SERS predicted residues could be involved in the interaction with DNA.

**Figure 13:**
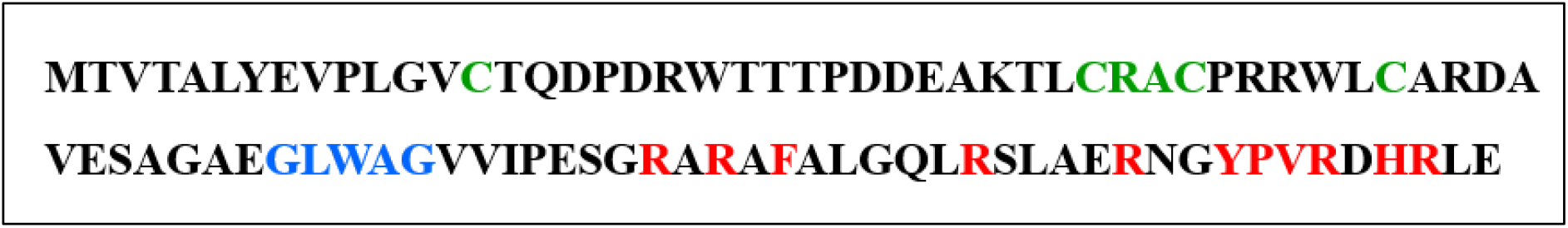
The sequence of the WhiB6 protein (obtained from the sequencing result of the WhiB6 plasmid). Residues highlighted in green are the conserved Cysteine residues, the amino acids in blue depict the conserved tryptophan region, and those in red are the residues predicted by SERS that were found to be involved in the WhiB6-*espA* interaction.

## CONCLUSION

We all have witnessed the health emergency caused by a global pandemic worldwide. We had seen how a virus had undergone various mutations and emerged into a new and more lethal variant. Hence, a disease-causing pathogen can change its form and become a contagious, life-threatening variant. *Mtb* is another fatal disease-causing bacterium that causes tuberculosis. This disease has always remained a massive problem and is still a threat to public health. The major problem encountered with this pathogen is its ability to gain resistance to the drugs used to cure the disease. Due to this reason, this disease always demands new drug targets and needs to find new medications to control the disease. Several proteins, genes, pathways, etc., of *Mtb*, have been targeted so far. Here, we have focused on targeting a crucial transcriptional regulator, i.e., the WhiB6 protein.

WhiB proteins are essential transcriptional regulators of *Mtb*. They can be a promising drug target to cure tuberculosis. However, detailed knowledge of the biophysical characteristics of the protein, its function, and how it regulates the transcription of other crucial genes has to be unveiled before using it as a drug target. Hence, we have focused on the WhiB6 protein, which regulates virulence and stress; we have characterized the protein and investigated the interaction of WhiB6 with the promoter DNA of the *espA* gene.

In the present study, the binding of *espA* promotor DNA with WhiB6 (a transcription factor) is explored using biophysical techniques such as fluorescence spectroscopy, CD spectroscopy, Raman spectroscopy, and ITC. The thermodynamic parameters obtained via ITC indicate that the binding of WhiB6 with DNA is a spontaneous (ΔG being negative) process. Moreover, a comparatively higher negative value of enthalpy and negative value of (TΔS) suggest more of an enthalpic-driven interaction which can be attributed to the formation of polar bonds, including hydrogen. Along with the thermodynamic feasibility, conformational changes are also observed in CD and steady-state fluorescence spectroscopy. It is observed that the secondary structure of WhiB6 is altered during the binding. There is a gradual decrease in the CD signal with increased DNA concentration. The SERS data illustrate that Arginine, proline, valine, phenylalanine, tryptophan, histidine, and tyrosine could be involved in the WhiB6-*espA* promoter DNA interaction. Furthermore, it is observed that MgCl_2_ enhanced the binding efficiency of interaction between WhiB6 and *espA* promoter DNA. Additionally, the oxidized apo-WhiB6 ((S)_2_) protein has a stronger binding affinity for DNA than the reduced state ((SH)_2_).

Genes regulated by WhiB6 execute their function only when they receive a signal from WhiB6 that binds to a particular region of the gene with a certain affinity. Hence, the present study’s findings will help synthesize/discover small molecules that could mimic the binding specificity and affinity of the WhiB6 protein and, therefore, artificially alter, activate, or deactivate the functions of the genes regulated by it. Thus, this will provide an initial foundation for proposing a suitable drug by exploring a series of drugs to inhibit similar interactions.

## SUPPLEMENTARY DATA

This article contains supplementary data.

## ACKNOWLEDGMENTS

S.K. would like to acknowledge the Indian Council of Medical Research (ICMR), the Government of India, for funding the research and providing the fellowship. We are grateful to Professor Bong-Jin Lee, Seoul National University of Korea, South Korea, for providing us with the plasmid of WhiB.

## AUTHOR CONTRIBUTIONS

S.D. supervised the research. S.D. and S.K. designed the experiments. S.K. carried out the experiments and performed data analysis. Teena carried out some additional experiments and analyzed the data. R.S performed Raman data acquisition and analysis under the supervision of S.S. S.K., and R.S are involved in writing the manuscript. S.D. and S.S. edited the manuscript.

## Abbreviation

TB: Tuberculosis
Mtb: Mycobacterium tuberculosis
CD: Circular dichroism
ITC: Isothermal titration calorimetry
SDS-PAGE: sodium dodecyl sulfate polyacrylamide gel electrophoresis

