## Supplementary Figures for "Biophysical Characterization and Interaction study of WhiB6 Protein of *Mycobacterium tuberculosis* with Nucleic Acid"

*__To whom correspondence should be addressed.__

Shashank Deep,

Professor,

Department of Chemistry

Indian Institute of Technology, Delhi (IIT Delhi)

Hauz Khas, New Delhi – 110016, INDIA

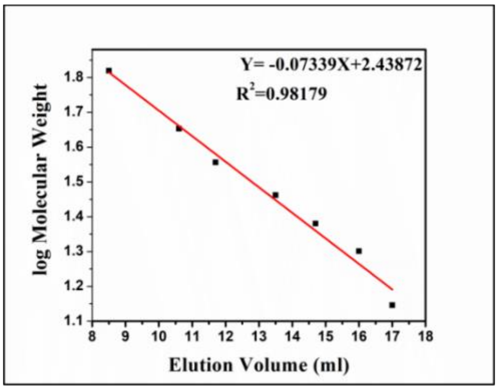


**Figure S1**: Calibration Curve plotted using standard proteins (for calibration of Superdex 200 Increase (10/300) GL column): α-Lactalbumin (14.2 kDa), Trypsin Inhibitor (20.1 kDa), Trypsinogen (24 kDa), Carbonic Anhydrase (29 kDa), Glyceraldehyde-3-Phosphate Dehydrogenase (36 kDa), Albumin, Egg (45 kDa), and Albumin, Bovine (66 kDa).

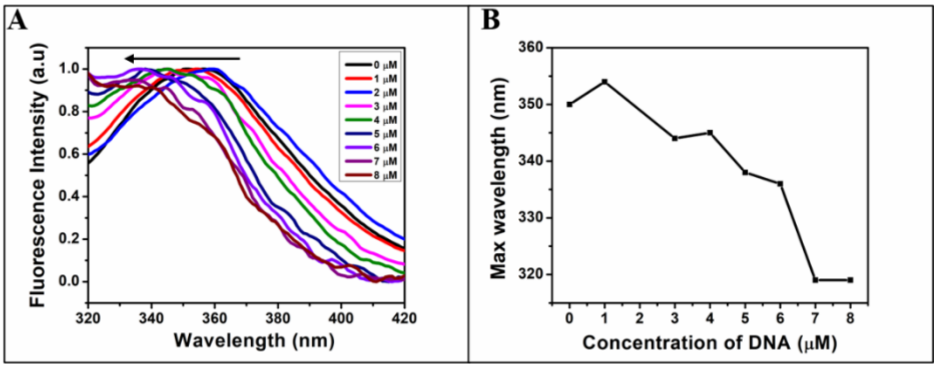


**Figure S2**: Normalized tryptophan emission spectra of WhiB6 showing a blue shift in the maximum wavelength upon interaction with espA promoter DNA (in the **absence of** $\mathbf{MgCl_2}$ in binding buffer). (**A**) Spectra showing a blue shift in the $\lambda_{max}$ (black arrow showing the direction of shift of the maximum wavelength). (**B**) plot showing the maximum emission wavelength at each concentration of DNA during the interaction.

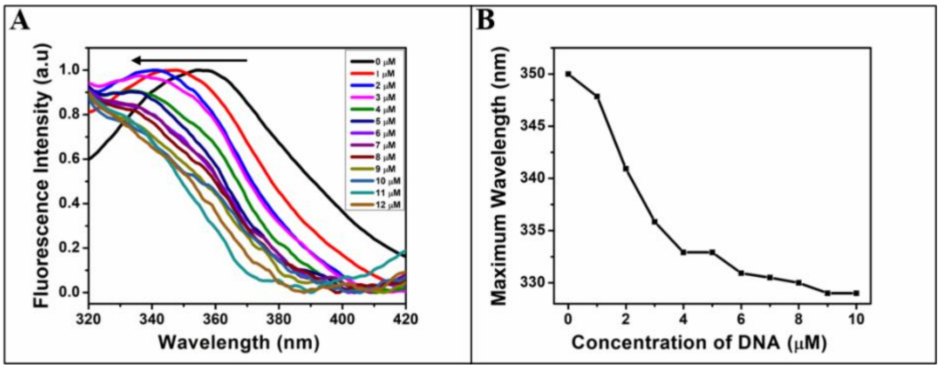


**Figure S3**: Normalized tryptophan emission spectra of WhiB6 showing a blue shift in the maximum wavelength upon interaction with *espA* promoter DNA (in the **presence of 10 mM** $\mathbf{MgCl_2}$ in binding buffer). (**A**) Spectra showing a blue shift in the $\lambda_{max}$ (black arrow showing the direction of shift of the maximum wavelength). (**B**) plot showing the maximum emission wavelength at each concentration of DNA during the interaction.

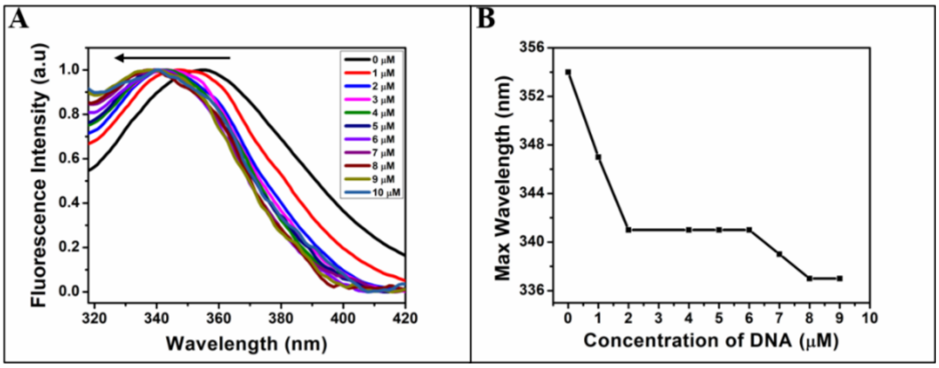


**Figure S4**: Normalized tryptophan emission spectra of WhiB6 showing a blue shift in the maximum wavelength upon interaction with *espA* promoter DNA (in the **presence of 10 mM $MgCl_2$** in binding buffer and WhiB6 is **reduced**). (**A**) Spectra showing a blue shift in the $\lambda_{max}$ (black arrow showing the direction of shift of the maximum wavelength). (**B**) plot showing the maximum emission wavelength at each concentration of DNA during the interaction.

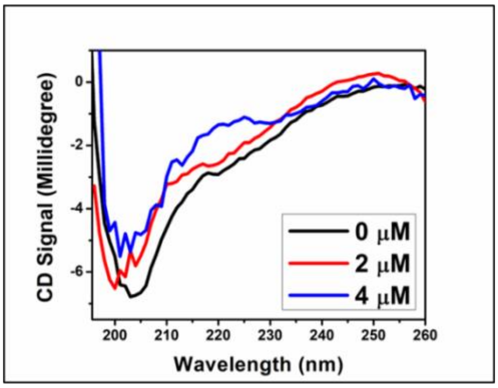


**Figure S5**: Far-UV Circular Dichroism spectra showing changes in the secondary structure of WhiB6 on interacting with *espA* promoter DNA (in the **absence of $MgCl_2$**).
